# Highly-connected, non-redundant microRNAs functional control in breast cancer molecular subtypes

**DOI:** 10.1101/652354

**Authors:** Guillermo de Anda-Jáuregui, Jesús Espinal-Enríquez, Enrique Hernández-Lemus

**Affiliations:** INSTITUTO NACIONAL DE MEDICINA GENOMICA

## Abstract

Transcriptional patterns are altered in breast cancer. These alterations capture the heterogeneity of breast cancer, leading to the emergence of molecular subtypes. Network biology approaches to study gene co-expression are able to capture the differences between breast cancer subtypes.

Network biology approaches may be extended to include other co-expression patterns, like those found between genes and non-coding RNA: such as microRNAs (miRs). Commodore miRs are microRNAs that, based on their connectivity and redundancy in co-expression networks, have been proposed as potential control elements of biological functions.

In this work, we reconstructed miR-gene co-expression networks for each breast cancer molecular subtype. We identified Commodore miRs in three out of four molecular subtypes. We found that in each subtype, each cdre-miR had a different set of associated genes, as well as a different set of associated biological functions. We used a systematic literature validation strategy, and identified that the associated biological functions to these cdre-miRs are *hallmarks of cancer*.

## Background

Breast cancer is a heterogeneous disease with many different manifestations. The heterogeneous nature of breast cancer can be observed at the transcriptional level, in the different gene expression patterns observed. This differences in breast cancer are at the basis of molecular classifications, such as the breast cancer molecular subtypes: Luminal A, Luminal B, Basal, and HER2-enriched. [22, 34]. These different molecular patterns are associated to different physiopathological properties, which can be used for clinical applications [8, 33].

The transcriptional patterns of breast cancer have been explored in previous works. Our group has found that representing the transcriptional program of breast cancer molecular subtypes as co-expression networks, it is possible by to capture the differences found between each cancer manifestation [15]. We have also shown how genes with coordinated expression patterns are found associated to each cancer subtype, and through these, it is possible to identify and associate functional perturbations to molecular subtypes [1, 2].

The regulatory programs of biological phenotypes are not limited to gene interactions. Elements such as non-coding RNAs are also involved in the regulation of gene expression. It has been shown that the transcriptional patterns of these non-coding RNAs also capture the heterogeneity of breast cancer molecular subtypes [31]. microRNA (miR) are a class of non-coding microRNAs that are currently a major study subject in cancer. Our group has developed work of studying these miRs from a network biology perspective [16].

Control in complex networks has important applications [26]. In the context of gene expression regulation, the control of gene expression, and more importantly, the concerted regulation of genes associated to biological functions, could have important biomedical applications. Similar concepts, such as master regulators[38, 30, 25], have been explored in different biological concepts, including cancer. In recent work, we introduced the concept of *Commodore miRs* (cdre-miRs): microRNAs that are highly connected and non-redundant in miR-gene co-expression netrowks in breast cancer, that are theoretically capable of controlling the state of specific biological functions by themselves [14]. In this work we intend to explore whether this *commodore* behavior can be found for miRs in networks of different breast cancer subtypes, how these cdre-miRs differ in each subtype, and how they are pontentially able to influence the activity of biological process important for the cancer manifestation.

## Materials & Methods

The workflow that was followed in this manuscript consists of the breast cancer gene and microRNA data acquisition, the co-expression network reconstruction, the identification of cdre-miRs, the functional enrichment of cdre-miR neighborhoods, and the literature validation of the identified biological functions. This workflow is represented in Figure 1.

**Figure 1:**
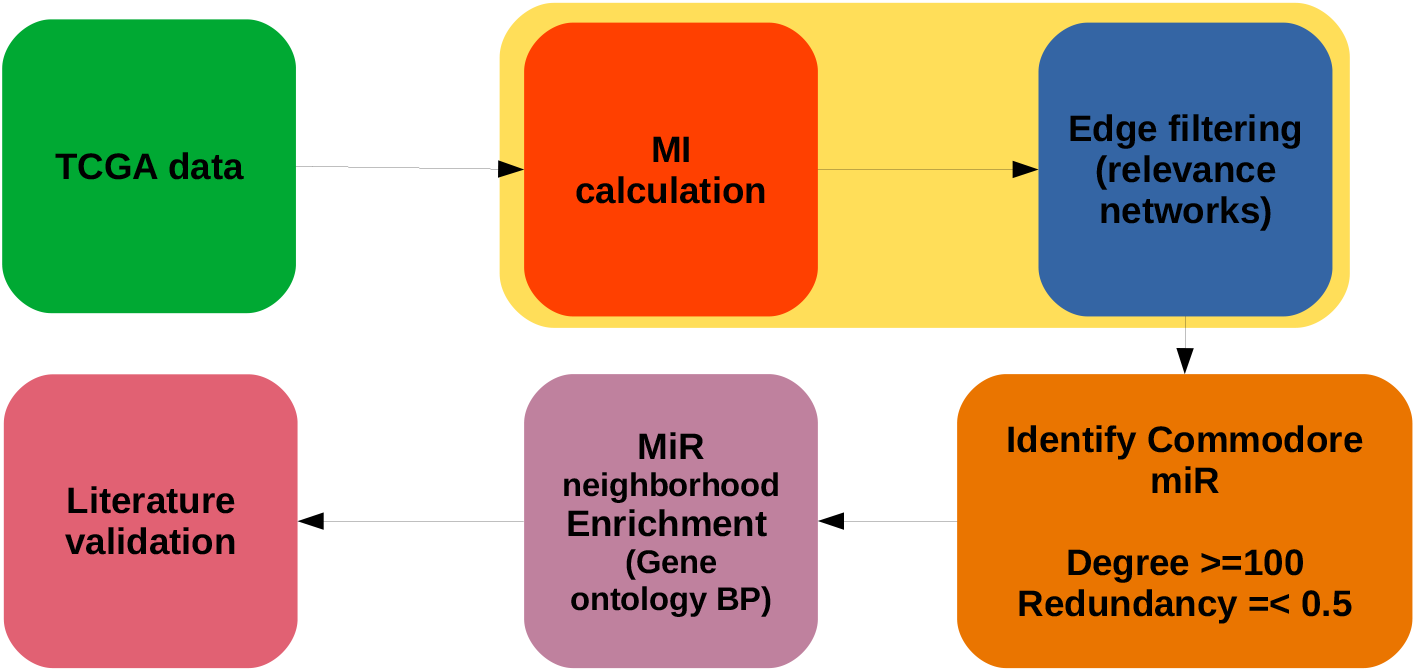
The cdre-miR analysis workflow

### Expression Data

Expression data for microRNA and genes in breast cancer was obtained from the Cancer Genome Atlas. The subset of breast cancer samples used in the 2012 TCGA publication [31] includes the molecular subtype sample classification. We acquired this information from the cBioportal website [6, 17]. We downloaded the expression data for gene and microRNA, for each molecular subtype: Luminal A (lumA), Luminal B (lumB), Basal, and HER2-enriched (HER2) from the Genome Data Commons website (https://portal.gdc.cancer.gov/repository).

The datasets found in the GDC platform are processed according to the bioinformatic pipelines found in https://docs.gdc.cancer.gov/Data/Bioinformatics_Pipelines/Expression_mRNA_Pipeline/ for genes and https://docs.gdc.cancer.gov/Data/Bioinformatics_Pipelines/miRNA_Pipeline/ for microRNA, which are referenced in the relevant original publications [31, 9]. For this work, we used FPKM - normalized data as expression values for mRNA, and RPMMM data as expression values for microRNA.

### microRNA/gene bipartite network reconstruction

We reconstructed a bipartite network representing the co-expression between microRNA and genes in each molecular subtype. For this, we used mutual information (MI) as a measure of miR-gene co-expression. Mutual information has been widely used for the reconstruction of co-expression networks [4, 28, 29, 42, 15, 7]. In the previous work by our group, we have successfully reconstructed miR-gene co-expression networks using this approach [16, 14].

For each molecular subtype, we calculate MI for each miR-gene pair based on their expression levels in order to fill an incidence matrix. We then select the miR-gene pairs that will be connected in the network based on their MI values. Those pairs with an MI value above a certain threshold are kept as links in the network, while those with an MI value below the threshold are discarded.

This strategy is the same that was used by our group in the previous miR-gene co-expression network manuscripts [16, 14]. The MI threshold selected for each network was set to be that which allowed us to keep the 0.9999 upper quantile of all possible links; this is based on a heuristic described by our group previously [12]. This allows to recover networks that have a comparable number of edges for each molecular subtype.

### Network analyses

The bipartite networks were analyzed for basic network topological properties using the igraph package for R [11]. The calculation of bipartite network properties, including the redundancy coefficient as defined in [24], were computed using the NetworkX package [19] for Python.

### Commodore miR identification

In our previous work regarding miR-gene co-expression networks [14], we defined the concept of a *Commodore miR* (cdre-miR): a microRNA that has a high number of neighbors, but a low redundancy coefficient (as defined by [24]) in a miR-gene co-expression network. In that work, we proposed considering a highly connected miR node to be, in the context of breast cancer, that which has a degree *k >*= 100, and a redundancy coefficient *rc* =*<* 0.5. In this work, we followed this definition.

### Functional enrichment of cdre-miR neighborhoods

Each identified cdre-miR has, by definition, a neighborhood of at least 100 genes. We identified biological functions that are associated to these neighborhoods, and therefore, to each cdre-miR. We performed this functional enrichment through an over-representation analysis, using the HTSanalyzer package for R [41]. We tested over-representation of the genesets encompassed in the Gene Ontology Biological Process (GO-BP) database [3, 10]. We considered a significance threshold of *Adjusted.p − value* =*<* 10^*−*3^ in the hypergeometric test.

### Functional category aggregation

We decided to present all the GO-BP categories found to be significantly associated to each cdre-miR. However, it is possible to leverage the ontology nature of the GO-BP database to group GO-BP categories that are both functionally related and composed of similar gene sets. To do this, we used the Wang similarity score [40], which measures the similarity between GO terms.

We calculated this similarity score for the GO-BP enriched for each cdre-miR of each subtype (using the GoSemSim package [43]). Then, we used this as the basis for a hierarchical clustering method, with which we generated for each cdre-miR, ten sets of functionally similar GO-BPs. We then selected as a *representative* GO-BP for each group, the GO-BP that had the lowest *Adjusted.p − value* within the group. The intention behind this is to obtain a more interpretable set of potential functional targets of cdre-miRs.

### Literature validation

We performed systematic queries to the Pubmed database to identify previously reported associations between the cdre-miRs and the functions identified in this work. To do so, we used the Rentrez package for R https://github.com/ropensci/rentrez. For each subtype, for each cdre-miR, we performed a query of the form *mir* + *RepresentativeGO* − *BP* considering each of the ten function groups associated to each cdre-miR.

## Results

### miR-gene co-expression networks

We reconstructed miR-gene co-expression networks for each molecular subtypes. These networks are comparable, by construction, in terms of the number of edges that they contain, and the number of miR and gene nodes (as they contain all the miRs and genes measured in the original experiments). The number of connected (*k >* 0) nodes and connected components (non-single nodes) in each network is variable, but they are, overall, comparable; this can be seen in Table 1. Visualizations of the largest connected components is found in the Figure 2. Other network parameters, including degree distribution, are found in Supplementary File 2.

**Table 1:** Network parameters

|  | Luminal A | Luminal B | Basal | HER2-enriched |
| --- | --- | --- | --- | --- |
| Connected nodes, miR | 269 | 384 | 414 | 587 |
| Connected nodes, gene | 2630 | 2731 | 2699 | 4011 |
| Edges | 6942 | 6942 | 6942 | 6951 |
| Connected components | 97 | 174 | 212 | 202 |

**Figure 2:**
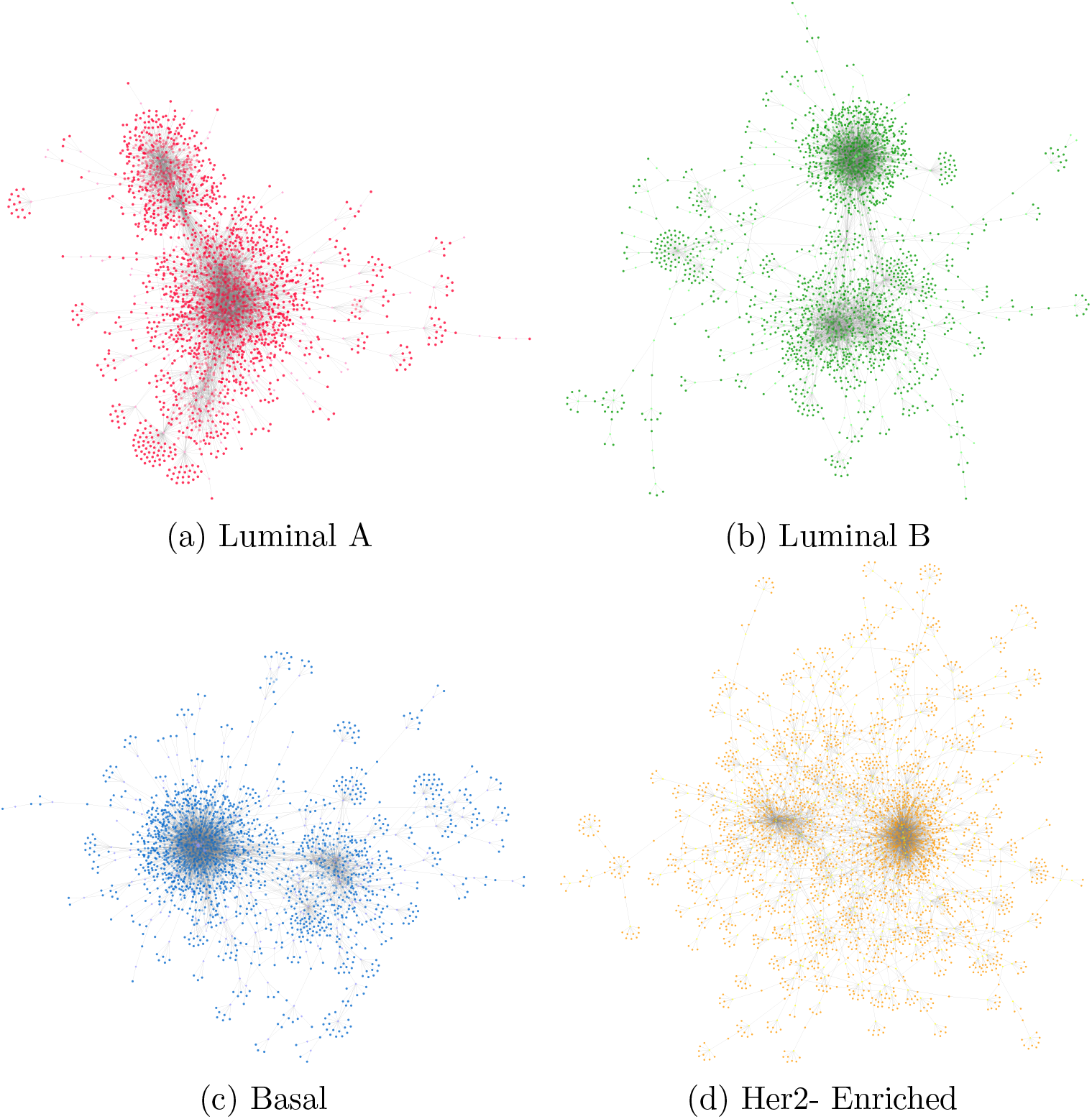
miR - gene co-expression network visualizations for each breast cancer molecular subtype; largest connected component shown.

It should be noted that in the case of HER2, we observe a slightly higher number of edges (6951 as opposed to 6942 in the rest of the subtype networks: this is explained due to the fact that there are links that have the exact same value than the threshold for HER2, and we did not implement any tie-breaking methods; we do not consider that the presence of these marginal edges may affect our downstream analyses.

### Identification of Commodore miRs: non-redundant, highly connected miRs

We identified 5 miRs that are non-redundant, and highly connected in at least one molecular subtype. These are:

- mir-139 and mir-150 in the Luminal A subtype
- mir-99a and mir-708 in the Luminal B subtype
- mir-136 and mir-139 in the Basal subtype

Figure 3 illustrates how these *commodores* are rare in the context of miRs in breast cancer subtypes. It should be noted that there are no cdre-miR in the HER2 molecular subtype, while each of the other subtypes possesses two cdre-miR.

**Figure 3:**
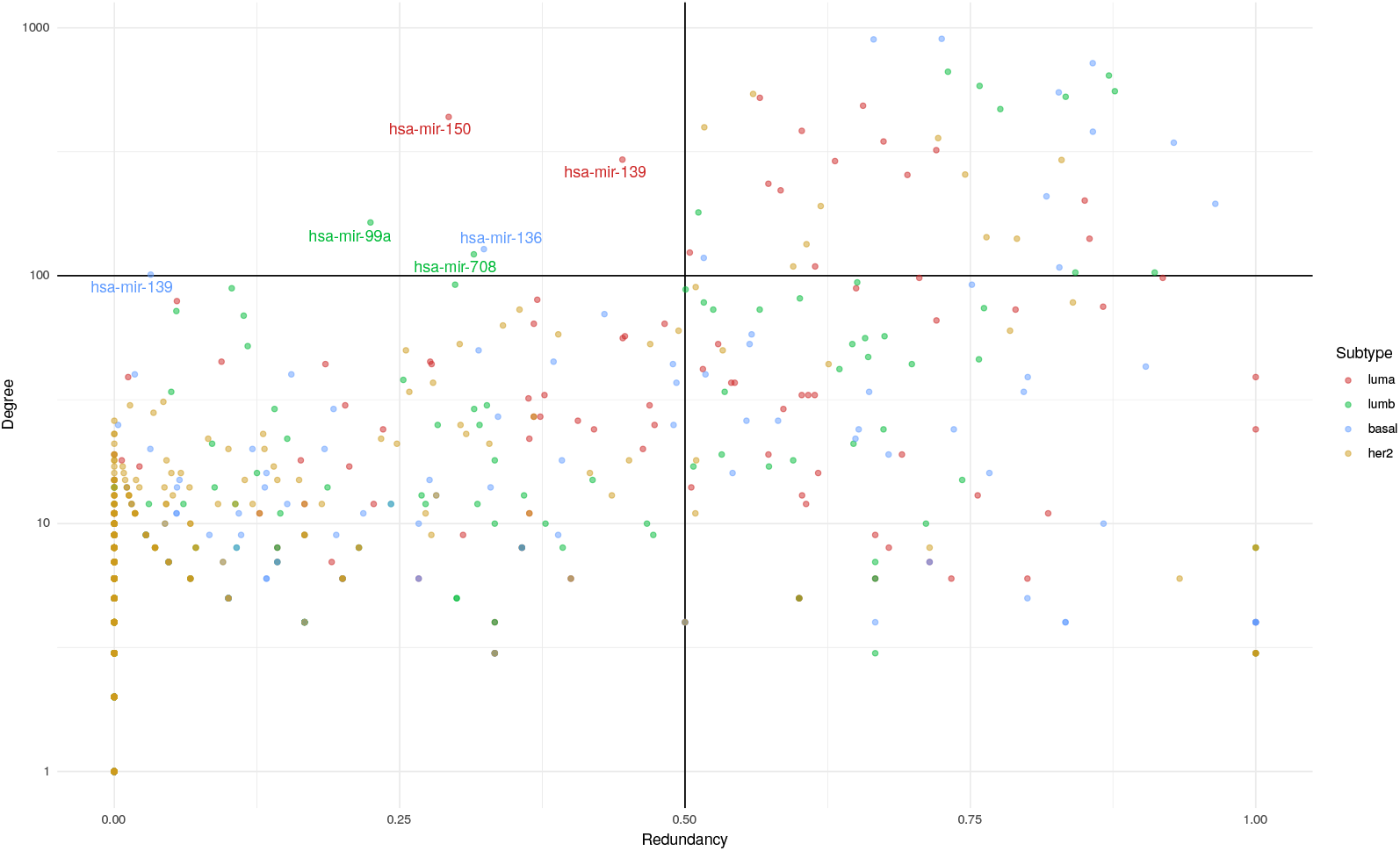
Scatter plot, degree vs redundancy coefficient for miR nodes in breast cancer molecular subtype networks. Each subtype is represented by a different color. The plot is divided by the commodore thresholds for degree (100) and redundancy coefficient (0.5). The upper-left quadrant contains *commodore miRs*.

Another thing to highlight is the fact that mir-139 is a commodore in both the Luminal A and Luminal B subtypes. The scatterplot shows, however, that they not exhibit the exact same behavior in terms of connectivity and redundancy. Supplemenary file 3 contains the degree and redundancy values of each cdre-miR in every subtype, which showcases that the behavior of miRs is different in each breast cancer manifestation.

### Functional enrichment of cdre-miR neighborhoods

We analyzed whether the neighborhoods of each cdre-miR could be associated to biological functions, by means of a hypergeometric test. We found that all cdre-miRs identified are linked in this fashion to a number of biological processes, as seen in Table 2. The whole set of enriched processes is found in Supplementary File 4.

**Table 2:** Enriched Gene Ontology Biological Processes in the gene neighborhoods of Commodore miRs

| subtype | miR | Enriched GO-BP |
| --- | --- | --- |
| Luminal A | hsa-mir-139 | 113 |
| Luminal A | hsa-mir-150 | 170 |
| Luminal B | hsa-mir-708 | 36 |
| Luminal B | hsa-mir-99a | 46 |
| Basal | hsa-mir-136 | 102 |
| Basal | hsa-mir-139 | 19 |

In Figure 4 we represent the biological processes associated to each cdre-miR as a network. This helps illustrate how there are some processes associated to several cdre-miR, while each cdre-miR has a set of processes that are uniquely associated to it. Since cdre-miRs are phenotype dependent, Figure 5 helps illustrate more clearly the way in which cdre-miRs are associated to different functions in each subtype. Finally, in panel5d the different behavior of mir-139 in the Luminal A and Basal subtypes is illustrated.

**Figure 4:**
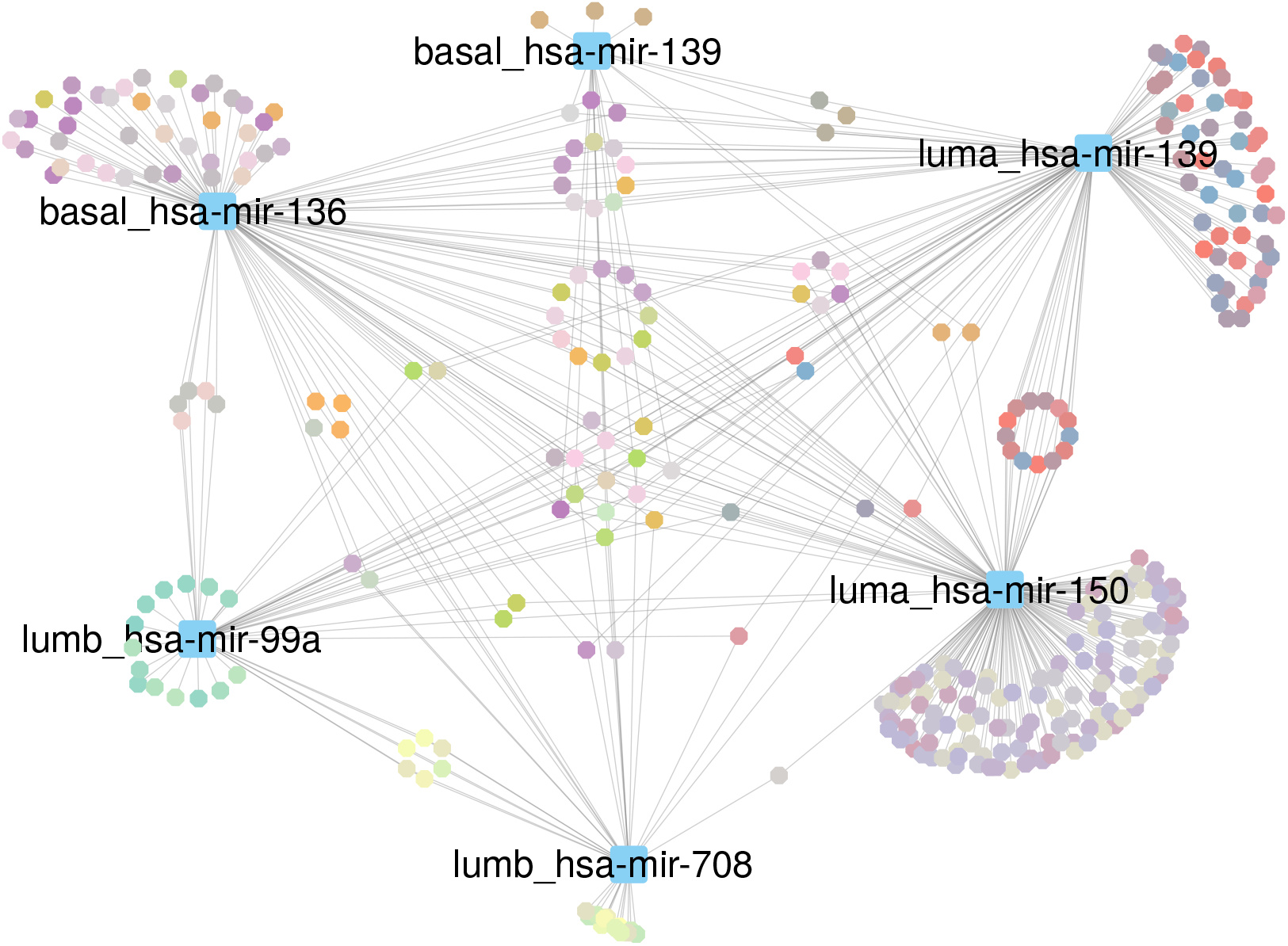
miR - Gene Ontology Biological Process network, containing all Commodore miRs found in each subtype. The color of GO-BP nodes represents groups of functionally similar processes.

**Figure 5:**
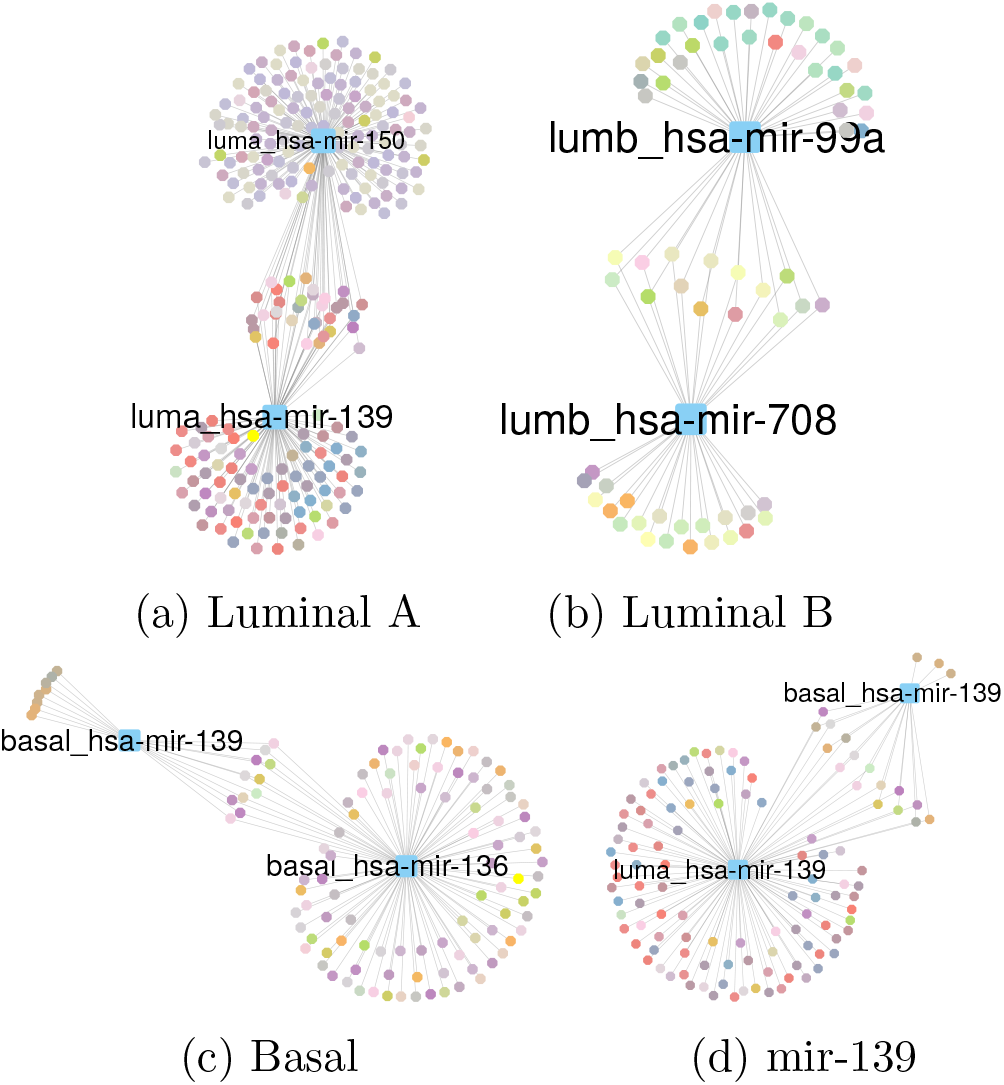
Commodore miR - Gene Ontology Biological Process for molecular subtypes (A-C) and the processes controlled by miR-139 in luminal A and basal subtypes. GO-BP node colors represent functionally similar processes.

Each panel in Figure 5 shows an overlap between the functions associated to each cdre-miR: this could be due to similarity between their respective neighborhoods. We provide, in Supplementary File 5, a similarity matrix of each cdre-miR neighborhood to show that this is not the case. In other words, each cdre-miR is affecting functions through different co-expressed gene sets.

### Biological processes aggregated by functional similarity

We grouped biological processes associated to each cdre-miR based on their functional similarity, as described in the methods section. The purpose of this was to reduce the number of GO-BP terms and aggregate them into the most representative (and biologically informative) terms. In Table 3 we show, for demonstration purposes, the characteristic terms for mir-139 in the Luminal A subtype. The full set of groups is provided as Supplementary File 6.

**Table 3:** Luminal A, miR-139

| group number | characteristic GO-BP | characteristic GO-BP name | Number of GO-BPs |
| --- | --- | --- | --- |
| 1 | GO:0001525 | angiogenesis | 14 |
| 2 | GO:0007186 | G protein-coupled receptor signaling pathway | 15 |
| 3 | GO:0010628 | positive regulation of gene expression | 7 |
| 4 | GO:0043066 | negative regulation of apoptotic process | 17 |
| 5 | GO:0006954 | inflammatory response | 15 |
| 6 | GO:0006936 | muscle contraction | 11 |
| 7 | GO:0006869 | lipid transport | 13 |
| 8 | GO:0006069 | ethanol oxidation | 9 |
| 9 | GO:0070374 | positive regulation of ERK1 and ERK2 cascade | 7 |
| 10 | GO:0098609 | cell-cell adhesion | 5 |

### Literature validation results

We systematically searched the biomedical literature to identify previous mentions of the identified biological functions associated to each cdre-miR. Table 4 shows the cdre-miR/function pairs for which at least one literature mention were found.

**Table 4:** Literature validation of biological functions associated to cdre-miRs

| Subtype | miR | GO representative term | Pubmed mentions |
| --- | --- | --- | --- |
| Luminal A | hsa-mir-139 | angiogenesis | 3 |
| Basal | hsa-mir-139 | angiogenesis | 3 |
| Luminal B | hsa-mir-708 | cell adhesion | 1 |
| Basal | hsa-mir-136 | cell adhesion | 5 |
| Basal | hsa-mir-139 | cell adhesion | 2 |
| Luminal A | hsa-mir-139 | negative regulation of apoptotic processes | 2 |
| Luminal A | hsa-mir-139 | positive regulation of gene expression | 7 |
| Basal | hsa-mir-136 | regulation of signaling receptor activity | 1 |
| Luminal A | hsa-mir-150 | signal transduction | 30 |

## Discussion

In previous work [12], we identified non-redundant, highly connected microRNAs in miR-gene co-expression networks of breast cancer. We proposed that this so-called “commodore” microRNAs are important regulatory elements, as they are potentially able to influence the expression level of a large set of genes by themselves; Furthermore, through this regulatory action, these microRNAs could be able to regulate specific biological processes. As such miR behavior was not found in healthy breast tissue networks, we speculated that cdre-miR could confer adaptational advantages to the tumor phenotype.

In this work, we explored cdre-miRs in the context of different manifestations of breast cancer: the molecular subtypes. We compared and contrasted these cdre-miRs, as well as their associated functions, and identified common and unique traits across the breast cancer landscape.

### Differences in microRNA roles in breast cancer sub-types

Considering that expression patterns are different between the molecular sub-types, we expected to find different sets of cdre-miRs associated to each molecular subtype. This was the case for three subtypes: Luminal A, Luminal B, and Basal. In the case of the HER2-enriched, we did not find any miR that was considered a commodore by our previously established definition.

For each of the remaining subtypes, we identified two miRs that we considered to be highly connected and non-redundant. None of the subtypes had the same pair of cdre-miR. Indeed, the only miR that had a commodore behavior in two subtypes was mir-139, in the Luminal A and Basal subtypes; nevertheless, as we mentioned in the Results section, this microRNA is linked to a different gene set, as well as a different function set, in each subtype.

Previous studies have shown that the expression patterns of molecular subtypes are different not only for genes, but also for microRNAs [31]. Since co-expression networks can be thought to be abstractions of the regulatory program behind these expression patterns [13], it is expected to find differences in these networks between subtypes: including differences in central nodes in the network. The fact that non-redundant, highly connected miRs emerge in most (but not all) subtypes could be indicative that having such regulatory element provides an advantage for the cancer phenotype.

### Possible advantages of commodore microRNAs for breast cancer

It is well known that microRNAs provide a regulatory mechanism for the control of gene expression [5]. Also known is the fact that microRNAs are widely derregulated in most cancer, although whether these are at the genesis of the disease, or a consequence of the pathological state, it is not known [27]. Since cancer is a complex disease, it is possible that both situations could happen, and even co-exist.

Potentially oncogenic microRNAs are able to confer functional features to cancer through their action as regulatory elements of gene expression [32]. Highly central microRNA nodes in miR-gene co-expression networks could act as control elements of gene expression based on their network connectivity, just like other gene elements have been identified [39]. By controlling the expression of genes involved in biological functions, these miRs could in turn control the activity of the function itself. In this context, commodore-miRs, both highly connected and non-redundant, could theoretically be the primary drivers of specific alterations of biological function.

### Functional heterogeneity and functional convergence

Having different cdre-miRs in each subtype leads to a varied landscape of altered functions. As we have shown, each cdre-miR in each subtype is associated to the expression of different genes, which in turn leads to differences in the associated functions. We observe that each cdre-miR has a set of functions that are unique to it, in the context of the phenotype in which it acts as a commodore. This could be one of the origins of the functional diversity observed and widely reported in breast cancer [18].

On the other hand, we observe that some functions may be affected by different cdre-miRs, either in the same or in different subtypes. The first explanation for this could be related to the (small) overlaps in gene neighborhoods observed between the cdre-miRs. But on a deeper level, this could be indicative of a convergence in biological process (de-)regulation; in other words, the control of a given function (or a significative subset of said function) confers an advantage to the tumor phenotype, which emerges regardless of the clinical (or molecular) manifestation, through different regulatory mechanisms. This could also be related to the lack of cdre-miRs in the HER2-enriched molecular subtype: being mostly driven by the amplification of a genomic region [23], the emergence of cdre-miRs is not needed for the development of this disease manifestation.

When we observe the terms that define the groups of functions associated to our cdre-miRs, it is can be observed that several of these refer to well known processes altered in cancer. Furthermore, when we look at the list of processes that were previously mentioned in the literature as being associated to breast cancer, we see that all of these belong to the set of functions known as the *hallmarks of cancer* [20, 21]. While experimental validation is still needed, if commodore-miRs are indeed acting as functional control elements specific to different breast cancer manifestations, then these could be attractive therapeutic options in the context of precision medicine [36, 37, 35].

## Conclusions

In this work we identify highly-connected, non-redundant (*commodore*) microRNAs in the context of breast cancer molecular subtypes. We found that different molecular subtypes exhibit different sets of these cdre-miRs, each associated to a specific set of biological functions. We observe that some of the associated functions are unique to each subtype, reflecting their functional diversity, while others are common. We found evidence in the literature that some of these functions being affected by our identified microRNAs; these functions are well known hallmarks of cancer, which could make targeting these microRNA a potential therapeutic alternative for different breast cancer manifestations.

## Supplementary files

Supplementary file 1: Mutual Information thresholds.

Supplementary file 2: Network parameters for each breast cancer molecular subtype.

Supplementary file 3: Degree and redundancy coefficient of commodore-miRs in all subtypes.

Supplementary file 4: Enrichment results for each cdre-miR neighborhood found in each subtype.

Supplementary file 5: Similarity matrix of cdre-miR neighborhoods.

Supplementary file 6: Functional category aggregation.

Supplementary file 7: Bipartite Networks for each molecular subtype.

Code for this manuscript can be found at https://github.com/guillermodeandajauregui/cdre-miR-BrCanSub

