## Supplementary File 1 for "Highly-connected, non-redundant microRNAs functional control in breast cancer molecular subtypes"

Supplementary File 1: MI threshold values corresponding to the 0.9999 quantile

| | MI threshold |
|---|---|
| Luminal A | 0.1807953 |
| Luminal B | 0.1606215 |
| Basal | 0.2086201 |
| HER2-enriched | 0.2121218 |
