## Supplementary File 2 for "Highly-connected, non-redundant microRNAs functional control in breast cancer molecular subtypes"

Supplementary File 2: Network Parameters

| | Luminal A | Luminal B | Basal | HER2-enriched |
|---|---|---|---|---|
| Nodes, miR | 1881 | 1881 | 1881 | 1881 |
| Nodes, gene | 36902 | 36902 | 36902 | 36902 |
| edges | 6942 | 6942 | 6942 | 6951 |
| Mir (degree !=0) | 269 | 384 | 414 | 587 |
| gene (degree != 0) | 2630 | 2731 | 2699 | 4011 |
| components | 35981 | 35842 | 35882 | 34387 |
| Components != 1 | 97 | 174 | 212 | 202 |
| largest component | 2494 | 2579 | 2470 | 3596 |
| miR, largest component | 156 | 194 | 190 | 362 |
| gene, largest component | 2338 | 2385 | 2280 | 3234 |

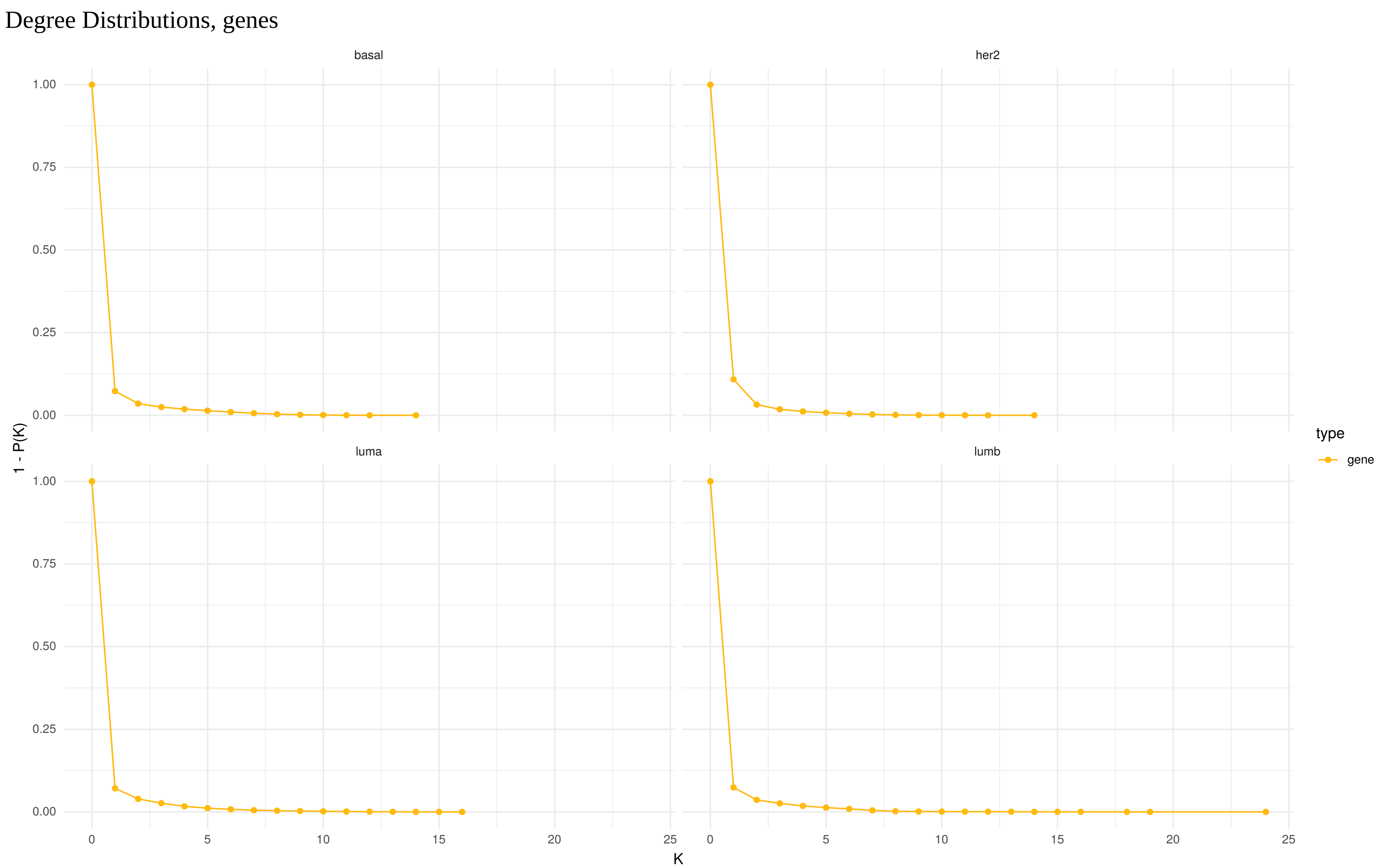

Degree Distributions, genes
basal
her2
luma
lumb
1 - P(K)
K
type
gene
1.00
0.75
0.50
0.25
0.00
0
5
10
15
20
25

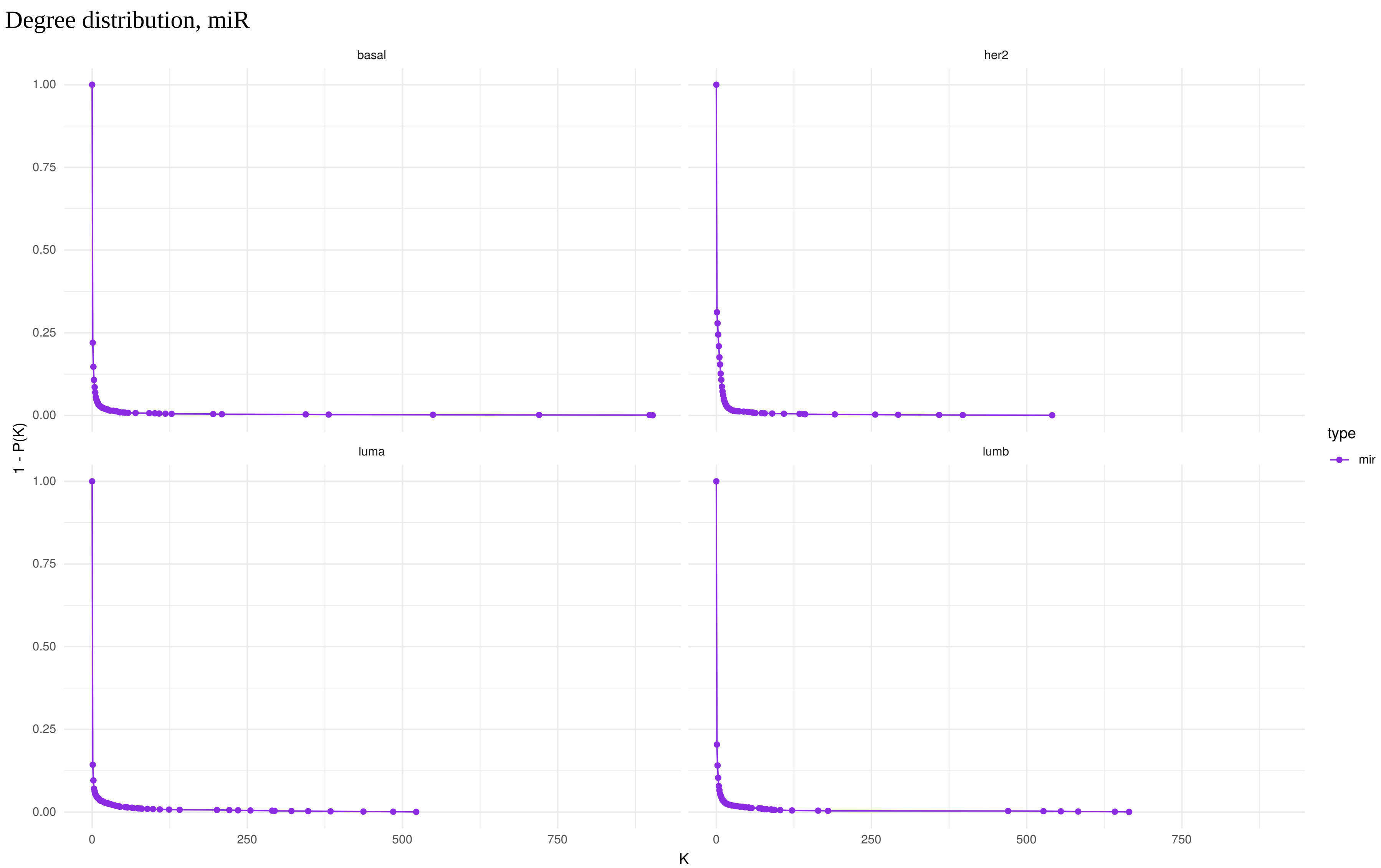

Degree distribution, miR
basal
her2
luma
lumb
1 - P(K)
K
1.00
0.75
0.50
0.25
0.00
0
250
500
750
type
mir
