## Supplementary File 3 for "Highly-connected, non-redundant microRNAs functional control in breast cancer molecular subtypes"

Supplementary File 3: Degree and Redundancy coefficient for each miRs with commodore behavior in at least one subtype. Values are highlighted in purple in the subtypes in which the miR is a cdre-miR.

| name | subtype | degree | Redundancy |
|---|---|---|---|
| hsa-mir-136 | Luminal A | 2 | 0.0000 |
| | Luminal B | 53 | 0.6466 |
| | **Basal** | **128** | **0.3238** |
| | HER2-enriched | 6 | 0.0000 |
| hsa-mir-139 | **Luminal A** | **294** | **0.4452** |
| | Luminal B | 52 | 0.1169 |
| | **Basal** | **101** | **0.0319** |
| | HER2-enriched | 18 | 0.0000 |
| hsa-mir-150 | **Luminal A** | **437** | **0.2930** |
| | Luminal B | 470 | 0.7762 |
| | Basal | 898 | 0.6652 |
| | HER2-enriched | 397 | 0.5169 |
| hsa-mir-708 | Luminal A | 1 | 0.0000 |
| | **Luminal B** | **122** | **0.3150** |
| | Basal | 13 | 0.2821 |
| | HER2-enriched | 8 | 0.0357 |
| hsa-mir-99a | Luminal A | 485 | 0.6560 |
| | **Luminal B** | **164** | **0.2245** |
| | Basal | 6 | 0.0000 |
| | HER2-enriched | 11 | 0.3636 |
